# PathoLive – Real-time pathogen identification from metagenomic Illumina datasets

**DOI:** 10.1101/402370

**Authors:** Simon H. Tausch, Tobias P. Loka, Jakob M. Schulze, Andreas Andrusch, Jeanette Klenner, Piotr W. Dabrowski, Martin S. Lindner, Andreas Nitsche, Bernhard Y. Renard

## Abstract

**Motivation:** Over the past years, NGS has become a crucial workhorse for open-view pathogen diagnostics. Yet, long turnaround times result from using massively parallel high-throughput technologies as the analysis can only be performed after sequencing has finished. The interpretation of results can further be challenged by contaminations, clinically irrelevant sequences, and the sheer amount and complexity of the data.

**Results:** We implemented PathoLive, a real-time diagnostics pipeline for the detection of pathogens from clinical samples hours before sequencing has finished. Based on real-time alignment with HiLive2, mappings are scored with respect to common contaminations, low-entropy areas, and sequences of widespread, non-pathogenic organisms. The results are visualized using an interactive taxonomic tree that provides an easily interpretable overview of the relevance of hits. For a human plasma sample that was spiked in vitro with six pathogenic viruses, all agents were clearly detected after only 40 of 200 sequencing cycles. For a real-world sample from Sudan the results correctly indicated the presence of Crimean-Congo hemorrhagic Fever Virus. In a second real-world dataset from the 2019 SARS-CoV-2 outbreak in Wuhan, we found the presence of a SARS Coronavirus as the most relevant hit without the novel virus reference genome being included in the database. For all samples, clinically irrelevant hits were correctly de-emphasized. Our approach is valuable to obtain fast and accurate NGS-based pathogen identifications and correctly prioritize and visualize them based on their clinical significance.

**Availability:** PathoLive is open source and available on GitLab (https://gitlab.com/rkibioinformatics/PathoLive) and BioConda (conda install –c bioconda patholive).

## 1 Introduction

The identification of pathogens directly from patient samples is a major clinical need. While highly accurate pathogen detection methods such as polymerase chain reaction (PCR), cell culture, or amplicon sequencing exist, such routine procedures often fail to identify the underlying cause of a patient’s symptoms due to their targeted behavior (Breitwieser, et al., 2015; Bzhalava, et al., 2013; Greninger, et al., 2017; Salzberg, et al., 2016). As a complementary approach, metagenomics Next-Generation Sequencing (NGS) has been proposed as a valuable technique for clinical application. NGS facilitates the detection and characterization of pathogens without a priori knowledge about candi-date species. Further, it generates a sufficient amount of data to detect even lowly abundant pathogens without targeted amplification of specified sequences allowing for hypothesis-free diagnostic analysis.

Current tools to address NGS-based pathogen identification can be divided into two major categories, either aiming to discover yet unknown genomes (Dutilh, et al., 2012; Huson, et al., 2016; Kostic, et al., 2011; Li, et al., 2016; Norling, et al., 2016; Piro, et al., 2017; Roux, et al., 2014; Skewes-Cox, et al., 2014; Tausch, et al., 2015; Wommack, et al., 2012; Zhao, et al., 2017) or to detect known organisms in a sample (Bray, et al., 2016; Byrd, et al., 2014; Dadi, et al., 2017; Flygare, et al., 2016; Francis, et al., 2013; Hong, et al., 2014; Lee, et al., 2016; Lindner and Renard, 2013; Menzel, et al., 2016; Naccache, et al., 2014; Piro, et al., 2019; Piro, et al., 2016; Scheuch, et al., 2015; Truong, et al., 2015; Wood, et al., 2019; Wood and Salzberg, 2014; Zheng, et al., 2017). From an algorithmic perspective, a further distinction can be made between alignment-based methods, alignment-free methods or combinations of both. While alignment-free methods usually deliver faster results, alignment-based methods potentially allow for a more extensive characterization of the sample.

Regardless of the algorithmic approach, existing methods based on unbiased metagenomics NGS face various obstacles, especially concerning the ranking of the results according to their clinical relevance and the long overall turnaround time (Breitwieser, et al., 2017; Dutilh, et al., 2017; Frey, et al., 2014; Lecuit and Eloit, 2014; Lecuit and Eloit, 2015; Mokili, et al., 2012; Roux, et al., 2017; Snyder, et al., 2009). The lack of good ranking methods is based on the fact that the distinction of clinically relevant and irrelevant data is not trivial. First, the dominating part of the sequences in a patient sample usually originates from the host genome. Second, there are nucleic acids of various species that are usually of low clinical relevance such as endogenous retroviruses (ERV) or non-pathogenic bacteria which commonly colonize a person. For these reasons, the number of reads hinting towards a relevant pathogen can be as low as a handful of individual reads. To put it more generally, it is a widespread misconception to rely only on quantitative measures when ranking the importance of candidate hits as not the amount but the uncommonness of a species in a given sample may give critical indications on its relevance. Based on the premise that a large proportion of the produced reads may stem from the host genome, species irrelevant for diagnosis, or common contaminations, even highly accurate methods struggle with false positive hits potentially concealing the relevant results. This central problem is getting worse when considering that even microbial databases are contaminated with human sequences (Breitwieser, et al., 2019). Existing pipelines tackle this problem in different ways. One common strategy is to ignore sequences that occur in a reference database of host and contaminating sequences (Byrd, et al., 2014; Dutilh, et al., 2012; Flygare, et al., 2016; Naccache, et al., 2014; Wommack, et al., 2012; Zheng, et al., 2017). While facilitating cleaner results, this approach may lead to a premature rejection of relevant sequences and does not solve the problem of human contaminations in reference databases as those “derive primarily from high-copy human repeat regions, which themselves are not adequately represented in the current human reference genome” (Breitwieser, et al., 2019). Further, the definition of precise contamination databases proves rather difficult and has not yet been adequately solved. Thus deleting any results to gain a better overview comes at great risk of overlooking the true cause of an infection. A different strategy are intensity filters, as implemented e.g. in SLIMM (Dadi, et al., 2017), that disregard sequences with low genome coverages. As the author states, this step eliminates many genomes which introduces the risk of losing information that might be relevant in the following diagnostic process. This problem even intensifies for marker-gene based methods such as MetaPhlAn2 (Truong, et al., 2015), as large parts of the sequenced reads cannot be assigned due to the miniaturized reference database. While this may lead to a better ratio of seemingly relevant assigned reads to those from the background, it comes with the risk of disregarding relevant candidates.

Another fundamental problem of NGS-based pathogen identification approaches is the fact that sequencing and analysis is very time consuming. Even when considering the reduction of sequencing time in the last years, current mid- and high-throughput devices still have maximum runtimes of more than a day (NextSeq 550) and up to two (NovaSeq 6000) or three days (HiSeq X), respectively. The resulting turnaround times of two to four days including data processing and analysis are not short enough for many critical scenarios such as sepsis and infectious disease outbreaks. To obtain actionable results within an appropriate time frame it is crucial to reduce the time span from sample receipt to diagnosis. However, existing approaches to speed up NGS-based diagnostics come with significant disadvantages such as a highly reduced throughput and data quality (Quick, et al., 2015), massive reduction of analyzed reads or targets (Stranneheim, et al., 2014) or the need of specialized hardware that involves additional costs and relatively low flexibility to adapt the workflow to a given scenario (Miller, et al., 2015). An actual approach for taxonomic classification of NGS data during runtime of sequencing is implemented in LiveKraken, a real-time version of the well-known Kraken software (Tausch, et al., 2018). However, by not providing positional information in the results, a sequence-based ranking to determine the relevance of hits is not possible with this approach.

As a general complement to real-time analysis of short-read sequencing data, there are several promising studies for pathogen detection using the MinION handheld device which is particularly useful for field studies and produces longer reads of up to several hundred kilobase pairs. While allowing very fast throughput times, these devices yield only approximately a million reads with comparably low per-base qualities, limiting their areas of application to targeted sequencing so far (Cao, et al., 2016; Greninger, et al., 2015; Loose, et al., 2016; Quick, et al., 2015; Stewart and Watson, 2017).

Therefore, from today’s perspective, NGS is the only technology providing sufficient amount and quality of data for many applications in clinical diagnostics. The currently high turnaround times from sample arrival to final diagnosis make it necessary to develop efficient methods to generate, analyze, and understand large metagenomics datasets in an accurate and quick manner to pave the way for NGS as a standard tool for clinical diagnostics. This enforces NGS-based diagnostics workflows to generate and evaluate large numbers of reads to facilitate adequate sequencing depths while reducing the time span between sample receipt and diagnosis. To overcome the named obstacles, we present PathoLive, an NGS-based real-time pathogen detection tool. We present an innovative approach to handle the occurrence of common contaminations, background data and irrelevant species in a single step. To tackle the problem of long overall turnaround times, we based our novel approach on the real-time read mapper HiLive2 that enables the analysis of sequencing data while an Illumina sequencer is still running (Loka, et al., 2019). This enables PathoLive to perform nucleotide-level analysis based on NGS providing an open view and high accuracy in short turnaround times while generating an intuitive and interactive visualization of results that highlights organisms of high clinical significance.

## 2 Methods

### Implementation

Our workflow follows a different paradigm than other frameworks to tackle the existing problems, as shown in Fig. 1: (i) prepare informative, well defined reference databases, (ii) automatically define contaminating or non-pathogenic sequences beforehand, (iii) use HiLive2 for accurate real-time alignment of Illumina sequencing data, (iv) visualize the potential risk of candidate pathogens and present results in an intuitive, comprehensible manner. The details on the modules for each of these steps are provided in the following paragraphs:

**Fig. 1.**
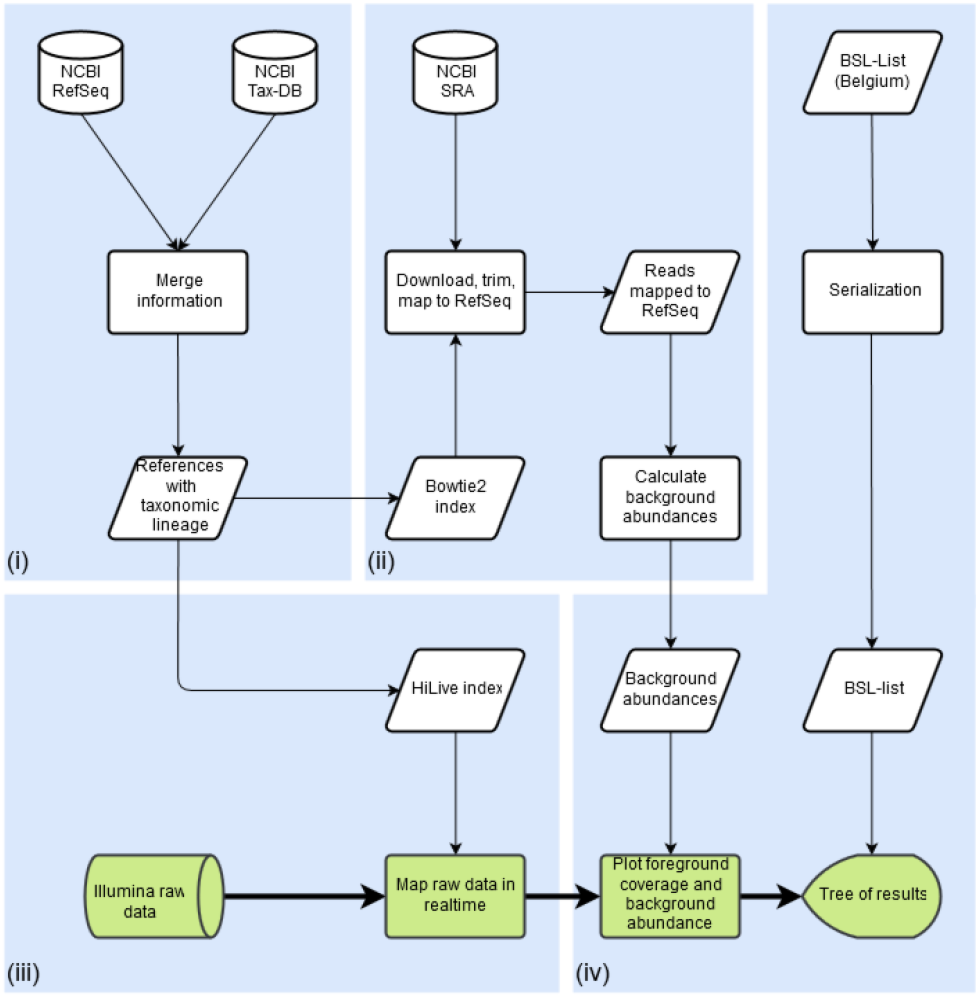
Workflow of PathoLive including four main modules. **(i)** Automated download and taxonomic tagging of reference information from NCBI RefSeq; **(ii)** NGS datasets from the 1000 Genomes Project are downloaded, trimmed and searched for database sequences from step (i), marking abundant stretches as clinically irrelevant; **(iii)** Reads from the clinical sample are mapped in real-time, producing intermediate alignment files; **(iv)** results are visualized in an easily understandable manner, providing all available information while pointing to the most relevant results. Only the steps highlighted in green are calculated in execution time, steps in white are precomputation. Graphical results are presented only minutes after the sequencer finishes a cycle if desired.

i. **Preparation of reference databases:** In order to save computational effort during the analysis, reference databases including the full taxonomic lineage of organisms are prepared before the first execution of PathoLive. For this purpose user selectable databases, for example the RefSeq Genomic Database (Brister, et al., 2015), are downloaded from the File Transfer Protocol (FTP) servers of the National Center for Biotechnology Information (NCBI) and annotated accordingly with taxonomic information from the NCBI Taxonomy Database. While preserving the original NCBI annotation of each sequence, additional information is appended to the sequence header. This information consists of each taxonomic identifier (TaxID), rank and name of each taxon in the lineage of an organism. Afterwards, user definable sub-databases of taxonomic clades relevant for a distinct pathogen search are automatically created. For the experiments in this manuscript, we focused on viruses. The database updater used for this purpose is available at https://gitlab.com/rki_bioinformatics/database-updater. The viral database used in this manuscript can be downloaded as a single compressed FASTA file from Zenodo (https://doi.org/10.5281/zenodo.2536788) and is ready to use for viral diagnostics with PathoLive.
ii. **Identification and labelling of clinically irrelevant hits**: A main obstacle in NGS based diagnostics is the large amount of background noise contained in the data. This includes various sources of contamination such as sequencing artefacts, ambiguous references and clinically irrelevant species, which hinder a quick evaluation of a dataset. Defining an exhaustive set of possible contaminations is a yet unachieved goal. Furthermore, deleting such sequences carries the risk of losing relevant results. Since in this step, raw sequencing data from a human host is examined, the logical conclusion is to contrast it to comparable raw datasets instead of processed genomes. We implemented a method to define and mark all kinds of undesired signals on the basis of comparable datasets from freely available resources. For this purpose, raw data from 236 randomly selected datasets from the 1000 Genomes Project Phase 3 (The 1000 Genomes Project Consortium, 2015) were downloaded, assuming that a large majority of the participants in the 1000 Genomes Project were not acutely ill with an infectious disease. The full list of selected datasets is provided in the supplementary material (Section 3.3). The reads are quality trimmed using Trimmomatic (Bolger, et al., 2014) and mapped to the selected pathogen reference database using Bowtie2 (Langmead and Salzberg, 2012). Whenever a stretch of a sequence is covered once or more in a dataset from the 1000 Genomes Project, the overall background coverage of these bases is increased by one. Coverage maps of all references from the pathogen database are stored in the serialized pickle file format. Stretches of DNA found in this data are marked as of lower clinical significance and visualized as such in later steps of the workflow. The coverage maps of the background abundances are plotted in red color against the coverage maps of the reads from the patient dataset in green color on the same reference (Fig. 2). This enables highlighting presumably relevant results without discarding other candidate pathogens, giving the researcher the best options to interpret the results in-depth but still in an efficient manner. The code for the generation of these databases is part of PathoLive.
iii. **Using HiLive2 for real-time alignment of reads:** We used HiLive2 (version 2.1) to produce real-time alignments of intermediate sequencing results. Thereby, the raw sequencing data is directly loaded in raw bcl file format without the need to perform a file conversion step. Alignments are updated with each new sequencing cycle and output in BAM format can be created for any sequencing cycle. As changes in the mapping positions mainly occur in early sequencing cycles, we recommend to create output in shorter intervals at the beginning of sequencing. Options for integrated demultiplexing and adapter trimming are available. For algorithmic details of HiLive2, we refer to Loka, et al. (2019).
iv. **Visualization and hazardousness classification:** A key hurdle in a rapid diagnostics workflow, which is often underestimated, is the presentation of results in an intuitive way. Many promising efforts have been made by different tools, e.g. providing coverage plots (Lindner and Renard, 2015; Naccache, et al., 2014) or interactive taxonomy explorers (Flygare, et al., 2016; Huson, et al., 2016). While being hard to measure and thus often ignored, the time it takes for groups of experts to assess the results and come to a correct conclusion should be considered. Our browser-based, interactive visualization is implemented in JavaScript using the data visualization library D3 (Bostock, et al., 2011). For an example of the visualization, see Fig. 4. While providing all available information on demand, the structure of a taxonomic tree allows an intuitive overview. Detailed measures are available on genus, family, species and sequence level. For the calculation of scores for a given node *n*, we define *t*(*n*) as the total number of read alignments to an underlying species of *n. b*(*n*) is the total number of bases being covered by all reads with respect to *n*. Accordingly, *b_bg_*(*n*) describes the number of bases being covered by the background database and *b_fg\bg_*(*n*) is the number of bases being covered by the foreground but not by the background data. In total, we provide three different scores for each node *n* of the tree:

a. Total Hits *T_n_*, representing the total number of hits to all underlying sequences in this branch: *T_n_*: = *t*(*n*)
b. Unambiguous Bases *U_n_*, representing the total number of bases covered in the foreground data but not in any background dataset: *U_n_*: = *b_fg\bg_*(*n*)
c. Weighted Score *W_n_*, being the ratio of unambiguous bases for the foreground data to the number of bases covered by the background database and logarithmically weighted by the total number of alignments: 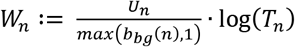

**Fig. 2.**
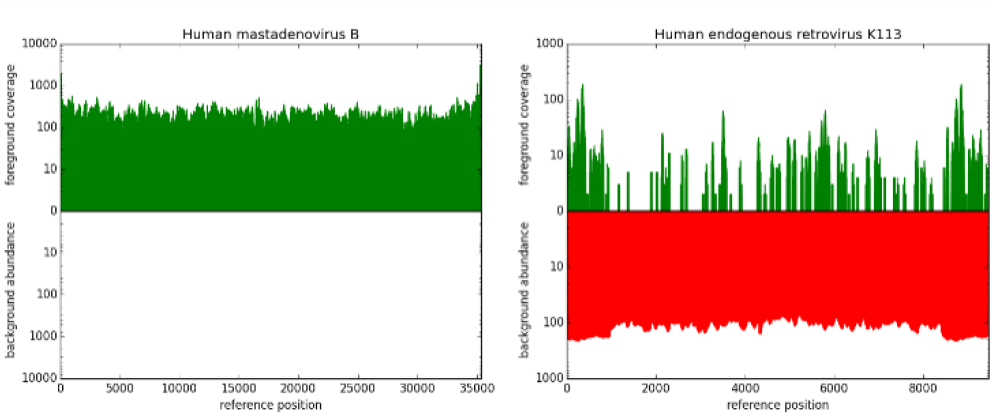
Two examples of fore- and background coverage plots. The upper, green bars show the coverage of a given genome in the foreground dataset, namely the reads sequenced from the patient sample. The lower, red part indicates in how many datasets from the 1000 Genomes Project a sequence is abundant. Bases covered in background datasets are regarded as less informative. Left: Fully covered genome of human mastadenovirus B, showing no hits resulting from data from the 1000 Genomes Project. Right: Coverage of human endogenous retrovirus (HERV) K113, partly covered in the patient dataset and completely covered in ~ll0 datasets from the 1000 Genomes Project. Based on these illustrations, Human mastadenovirus B can be considered a relevant hit while HERV K113 is rightly found in the dataset, but not considered a clinically relevant candidate due to its common prevalence in healthy human individuals.

While the Total Hits *T_n_* can be useful to get a general impression of the abundance of sequences in the sample, the Unambiguous Bases *U_n_* provides a first comparison to the background dataset. The Weighted Score *W_n_* introduces an intensified metric of how often a sequence is found in a healthy individual, and thereby allows drawing stricter conclusions from the background data. Not only exactly overlapping mappings of fore- and background are regarded, but also the overall abundance of a sequence within the background data is considered.

The values of the selected scoring scheme are reflected in the thickness of the branches, which draws the visual focus to higher rated branches. Users can switch between the three scores via the respective buttons in the interactive visualization. In order to enable users to make early decisions regarding the handling of a sample as well as to further enhance the intuitive understanding of the results, the hazardousness of detected pathogens is color-coded based on a Biosafety level (BSL) score list (Biosafety and Biotechnology Unit, 2008). To improve BSL classification, minor changes were manually applied to improve matches to the organism names in the reference database. The BSL score gives information on the biological risk emanating from an organism. Therefore, it qualifies as a measure of hazardousness in this use case. The BSL-score is color-coded in green (no information/BSL1), blue (BSL2), yellow (BSL3) or red (BSL4), and the maximum hazardousness-level of a branch is propagated to the parent nodes. Phages are displayed in grey, as they cannot infect humans directly, but may imply information on the presence of bacteria.

Details about the sums of all three available scores of all underlying species are provided on mouse-over (Fig. 4 in the results section). When expanding a branch to sequence level, additional plots of the foreground coverage calculated in step (iii) as well as the abundance of bases in the background datasets calculated in step (ii) are shown when hovering the mouse over the node (Fig. 2). These plots provide a visualization of the significance of a hit. The hits of a species in the patient dataset are shown in green, while background hits are drawn in red on a coverage plot. This way, it is easy to evaluate if a sequence is commonly found in non-ill humans and therefore can be considered less relevant, or if a detected sequence is unique and could lead to more certain conclusions.

### Validation

We compared the results of PathoLive to two existing solutions, Clinical Pathoscope (Byrd, et al., 2014) and Bracken (Lu, et al., 2017). We selected Clinical Pathoscope for its very sophisticated read reassignment method, which promises a highly reliable rating of candidate hits. It also is perfectly tailored to this use case. Other promising pipelines such as SURPI (Naccache, et al., 2014) or Taxonomer (Flygare, et al., 2016) were not locally installable and had to be disregarded. Bracken, a method based on metagenomics classification with Kraken (Wood and Salzberg, 2014), was included in the benchmark as one of the fastest and best known classification tools which makes it one of the primary go-to methods for many users. The experiment is based on a real sequencing run on an Illumina HiSeq 1500 in High Output Mode. We designed an in-house generated sample in order to have a solid ground truth. We ran all tools using 40 threads, starting each at the earliest possible time point when the data was available from the sequencer in the expected input format. For the non-real-time tools, the base calling was executed via Illumina’s standard tool bcl2fastq and the runtime was regarded in the overall turnaround time. Clinical Pathoscope and Bracken were both run with default parameters, apart from the multithreading. We built the databases for PathoLive and Bracken using the viral part of the NCBI RefSeq (O’Leary, et al., 2016). For Clinical Pathoscope we downloaded the associated database from http://www.bu.edu/jlab/wp-assets/databases.tar.gz using the provided viral database as foreground and the human database as background. Details of the database construction are given in the supplementary methods (Section 3.1). Please note that, in contrast to all other results shown in this manuscript, the live analysis of the in-house sample was performed using read-mapping results of HiLive, the predecessor of HiLive2. However, we repeated the analysis using HiLive2 and obtained similar results with respect to accuracy (cf. Supplementary Fig. 1 and Supplementary Table 1).

To validate PathoLive on real data, we applied it to a previously described diagnostic human serum sample from an outbreak of hemorrhagic fever virus in Sudan (Andrusch, et al., 2018; Kohl, et al., 2016) and a dataset from an outbreak of Severe acute respiratory syndrome coronavirus 2 (SARS-CoV-2) in Wuhan, 2019 (Wu, et al., 2020). As the data was only available in FASTQ format, it was converted to bcl file format following the procedure described in the supplementary methods (Section 3.2). The total read length was 2 x 301bp for the CCHFV dataset from Sudan and 2 x 151bp for the SARS-CoV-2 dataset from Wuhan.

### Sample preparation

Viral metagenomics studies were performed with a human plasma mix of six different RNA and DNA viruses as well-defined surrogate for clinical liquid specimen. The informed consent of the patient has been obtained. This 200 μL mix contained orthopoxvirus (Vaccinia virus VR-1536), flavivirus (yellow fever virus 17D vaccine), paramyxovirus (mumps virus vaccine), bunyavirus (rift valley fever virus MP12-vaccine), reovirus (T3/Bat/Germany/342/08) and adenovirus (human adenovirus 4) from cell culture supernatant at different concentrations. The sample also contains dependoparvovirus as proven via PCR. The sample was filtered through a 0.45 μM filter and nucleic acids were extracted using the QIAamp Ultrasense Kit (Qiagen) following the manufacturers’ instructions. The extract was treated with Turbo DNA (Life Technologies, Darmstadt, Germany). cDNA and double-stranded cDNA (ds-cDNA) synthesis were performed as previously described (70). The ds-cDNA was purified with the RNeasy MinElute Cleanup Kit (Qiagen). The purification method takes ~6h to complete. The Library preparation was performed with the Nextera XT DNA Sample Preparation Kit following the manufacturers’ instructions (Illumina). NGS libraries were quantified using the KAPA Library Quantification Kits for Illumina sequencing (Kapa Biosystems). If the starting amount of 1 ng of nucleic acid was not reached the entire sample volume was added to the library.

The diagnostic sample from Sudan was prepared according to (Andrusch, et al., 2018; Kohl, et al., 2016), including inactivation of the human serum in Qiagen Buffer AVL, extraction with Qiagen QIAamp Viral RNA Mini Kit and DNA digestion using the Thermo Fisher TURBO DNA-free Kit. A sequencing library was created using the Illumina Nextera XT DNA Library Preparation Kit. The sample was sequenced on an Illumina MiSeq.

The dataset from the outbreak of SARS-CoV-2 in Wuhan in 2019 was sequenced on an Illumina MiniSeq sequencing device and is publicly available at the NCBI Sequence Read Archive (SRA) under accession number SRR10971381 (Wu, et al., 2020).

## 3 Results

### 3.1 Pathogen detection in a spiked viral mixture

The human plasma sample spiked with a viral mixture was sequenced on an Illumina HiSeq 1500 in High Output mode on one lane. PathoLive was executed from the beginning of the sequencing run using 40 threads. Intermediary results were taken after 40, 60, 80 and 100 cycles or after 36, 55, 74 and 93 hours, respectively. The time needed to produce results from the intermediary sequencing data was lower than 25 minutes for all output cycles. Raw reads usable for the testing of other tools were available only after 95 hours as they had to be translated into the human readable fastq-format first. As a ground truth, we selected all sequences associated to the species described as abundant above.

The area under the curve (auc) of the receiver operating characteristic (ROC) was calculated using the 14 highest ranking species, as given by the tested tools. The top 14 of the identified species are considered because hits appearing after twice the number of true positives cannot be expected to be regarded by a user in this experiment. Furthermore, none of the tested tools found more true positives within the next 50 hits. The ROC-plot (Fig. 3) denotes the True Positive Rate and False Positive Rate for each threshold n≤14, whereby a threshold n means that the best n hits are taken into account. This means that only the rank of the hits was considered while disregarding the actual score. For PathoLive, the ranks were determined by the Weighted Score *W_n_*, for Clinical Pathoscope we used the “final guess” metric and for Bracken, the species with most estimated reads were ranked highest.

**Fig. 3.**
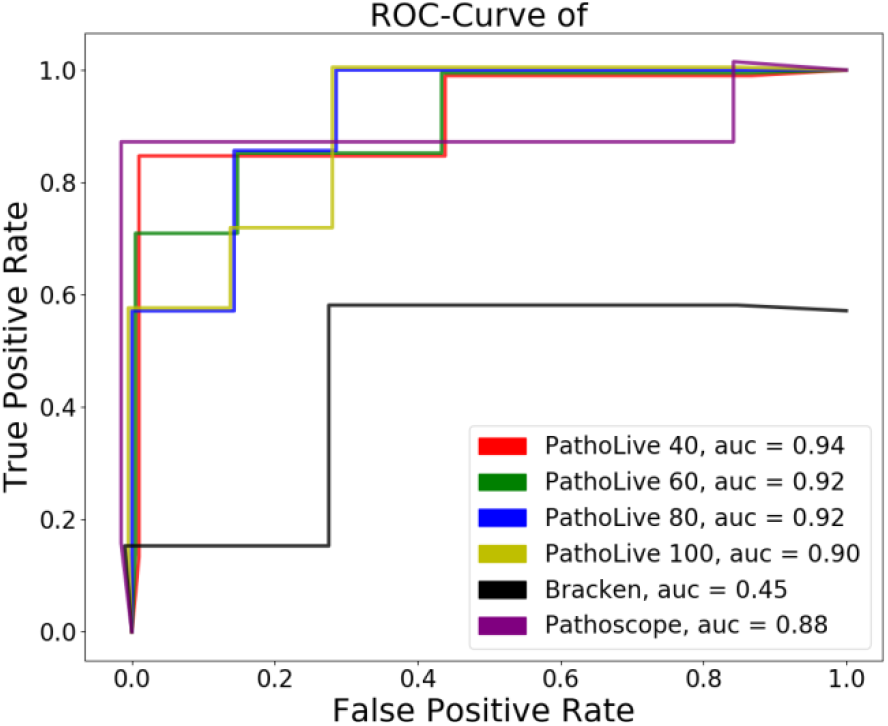
ROC-plot of benchmarked tools on a spiked dataset. Lines have slight offsets in x- and y-dimensions for reasons of distinguishability. We compared PathoLive to Clinical Pathoscope and Bracken on a human sample containing 7 viruses. PathoLive performs best regarding the ROC-auc at all sampled times (cycle 40, 60, 80 and 100) when compared to the results of the other tools after sequencing the complete first read

We were able to detect all abundant spiked species in the library after only 40 cycles of the sequencing run using PathoLive. While the overall number of false positive hits decreases with the sequencing time, the weighted score and the number of unambiguous bases yield accurate results throughout all reports. Reported phages are included in these numbers, although they are optically grayed out in the visualization, as they cannot infect vertebrates directly. As an example report, a screenshot of the resulting interactive tree of results after 80 cycles is shown in Fig. 4.

**Fig. 4.**
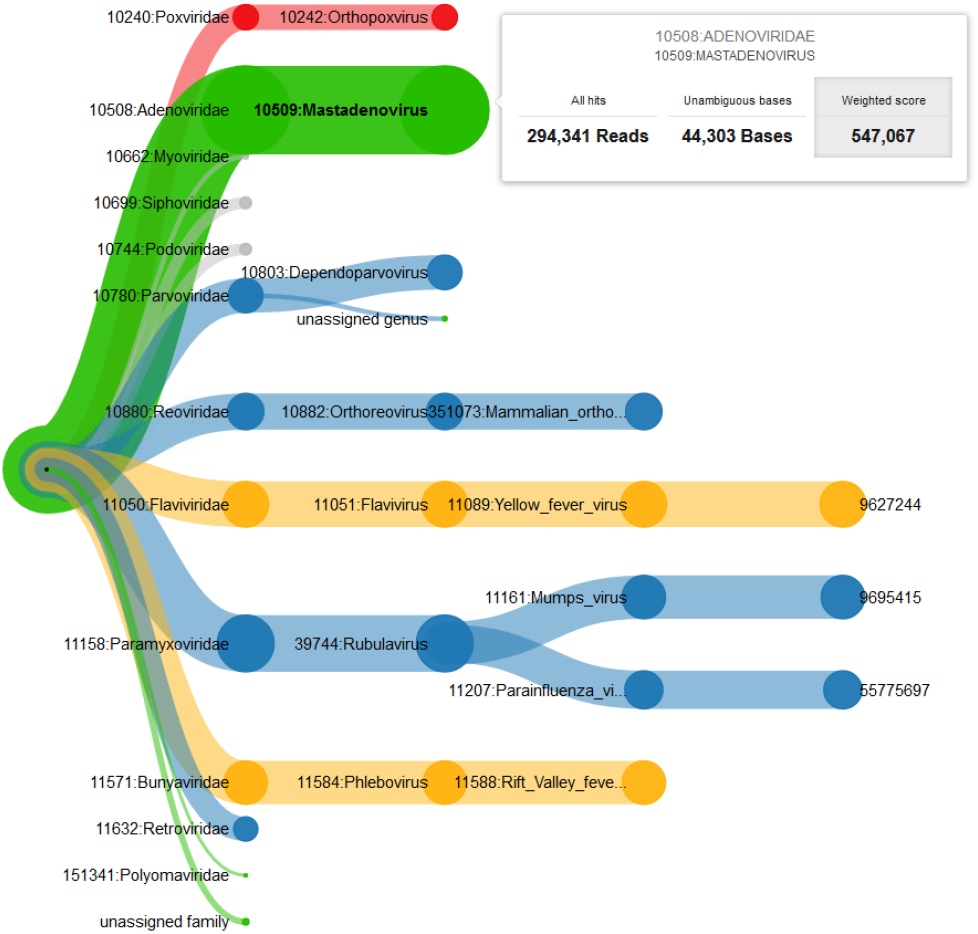
Example of the interactive taxonomic tree of results. The results show the described plasma sample at cycle 80 based on the weighted score. Thickness of the branches denotes the sum of scores of underlying sequences. The color codes for the maximum of the underlying BLS-levels (red=4, yellow=3, blue=2, green=1 or undefined; phages are shown in grey). On mouse-over, detailed information (here on genus Mastadenovirus) is displayed. The selected score (here: weighted score) is highlighted in grey. The visualization clearly emphasizes all spiked pathogens through the thickness of their clades, while other species are shown only in smaller clades and therefore ranked lower.

### 3.2 Identification of Crimean-Congo hemorrhagic Fever Virus in a real sample from Sudan

A central issue in pathogen identification, especially for viruses, is the potentially low number of pathogenic reads in the sample. Therefore, we demonstrated the performance of PathoLive on real data that is known to contain a low number of reads of interest. We analyzed a human serum sample from Sudan that was confirmed via PCR to contain Crimean-Congo hemorrhagic Fever Virus (CCHFV) but only shows a small amount of related reads in the corresponding Illumina sequencing data (45 out of 1,178,054 reads were reported by Andrusch et al. in 2018 (Andrusch, et al., 2018) to unambiguously belong to CCHFV). When running PathoLive with default parameters and having adapter trimming activated, Bunyaviridae was the family with the highest weighted score over the complete sequencing procedure when not considering phages and the “unassigned family” branch. Thereby, the score of Bunyaviridae was consistently equal to the score of the underlying species CCHFV while other underlying species didn’t contribute to the overall score of the family. Fig. 5 shows the development over time for all families that reach a score of 500 in at least one output cycle. It can be seen that the weighted score of CCHFV (represented by the family of Bunyaviridae) is in the top three of all identified families after only 30 sequencing cycles which corresponds to 5% of the sequencing procedure. At this time point, only 16 reads were aligned to CCHFV. Thus, indications for the correct finding are already possible within a short time span and based on only a couple of available reads while the result is more and more emphasized with ongoing sequencing. The only other family reaching a score higher than 500 and not exclusively containing phages was Retroviridae, being mainly driven by the species HIV1. However, a more detailed view on the sequence level shows that all mappings to HIV1 cluster in a small region of approximately 1,000bp (Fig. 6d) while the alignments to CCHFV distribute over the complete genome (Fig. 6b,c). This strongly indicates that CCHFV is more likely to be a true positive. Fig. 6 further shows the family level visualization of the PathoLive tree structure (Fig. 6a) and an example for Granulovirus of the Baculoviridae family that shows a high total number of mappings, but all of those being located in regions that are covered in the background database leading to a weighted score of 0 (Fig. 6e). The overall results for this sample show the strength of PathoLive to pronounce interesting findings at first glance while still allowing for a more detailed perspective that is often important for interpretation.

**Fig. 5.**
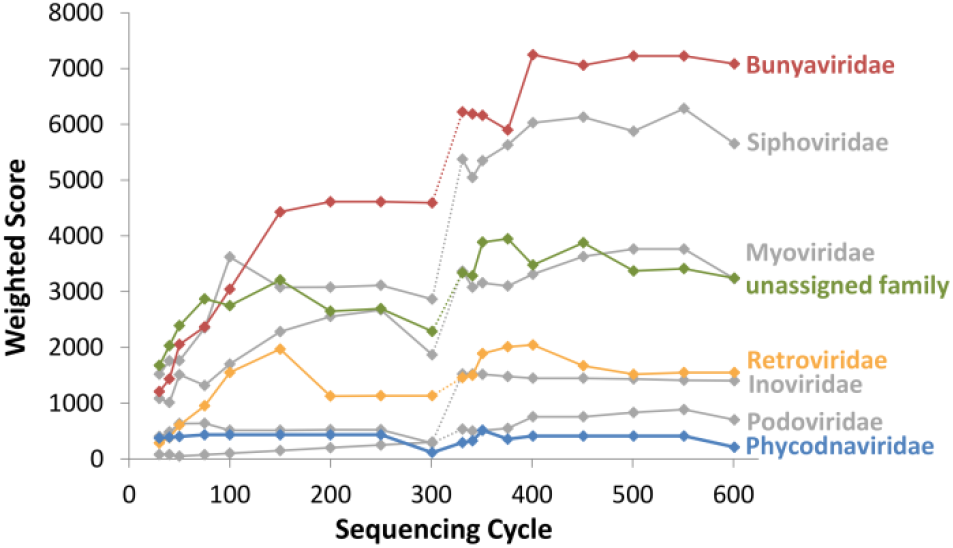
Development of the weighted score calculated by PathoLive over the sequencing procedure for all families reaching a score higher than 500 in at least one output cycle. Colors of the plots correspond to the underlying biosafety level in the last cycle, i.e. green for BSL-1, blue for BSL-2 and yellow for BSL-3. Phages are displayed in gray color. The dotted section of each line indicates the shift from the first to the second read of the 2×301bp data.

**Fig. 6.**
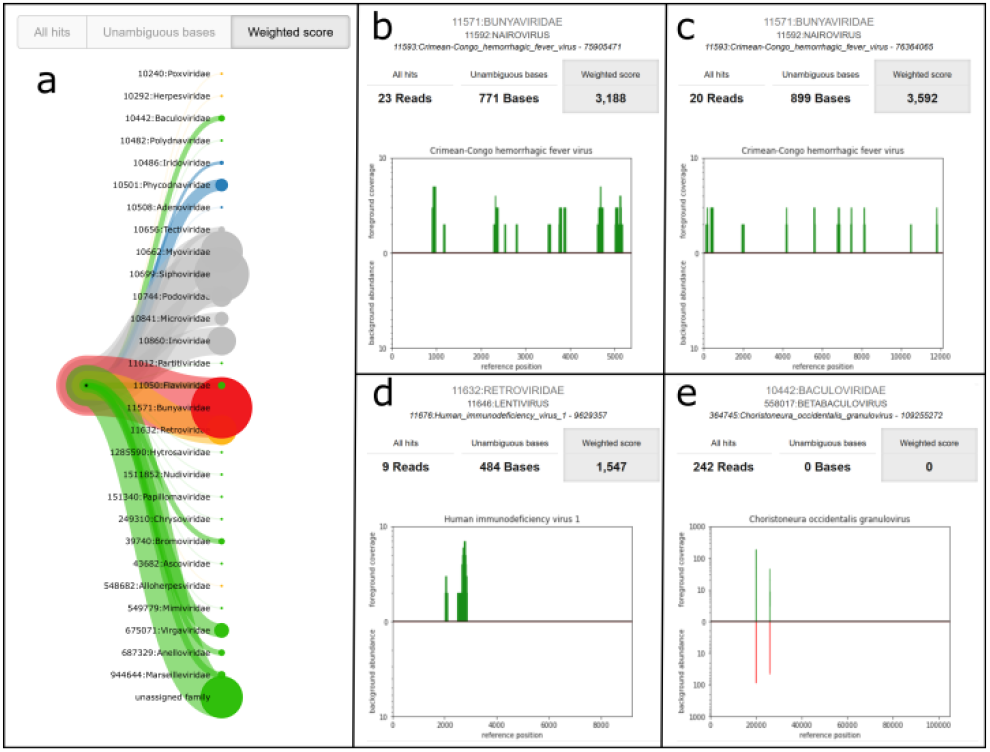
Visualization of the final results of PathoLive for cycle 602. (a) Tree structure on family level. (b,c) Tooltips for the sequence level of alignments for two CCHFV reference sequences of the Bunyaviridae family. (d) Tooltip for the sequence level of alignments for HIV1 of the Retroviridae family. (e) Tooltip for the sequence level of alignments for a Granulovirus reference of the Baculoviridae family.

### 3.3 Detection of a Coronavirus in a real sample from the 2019 SARS-CoV-2 outbreak in Wuhan

For a dataset from the outbreak of SARS-CoV-2 in Wuhan, 2019, we could also identify a Coronavirus as the most probable causative virus. This example clearly demonstrates the strength of our scoring approach.

**Fig. 7.**
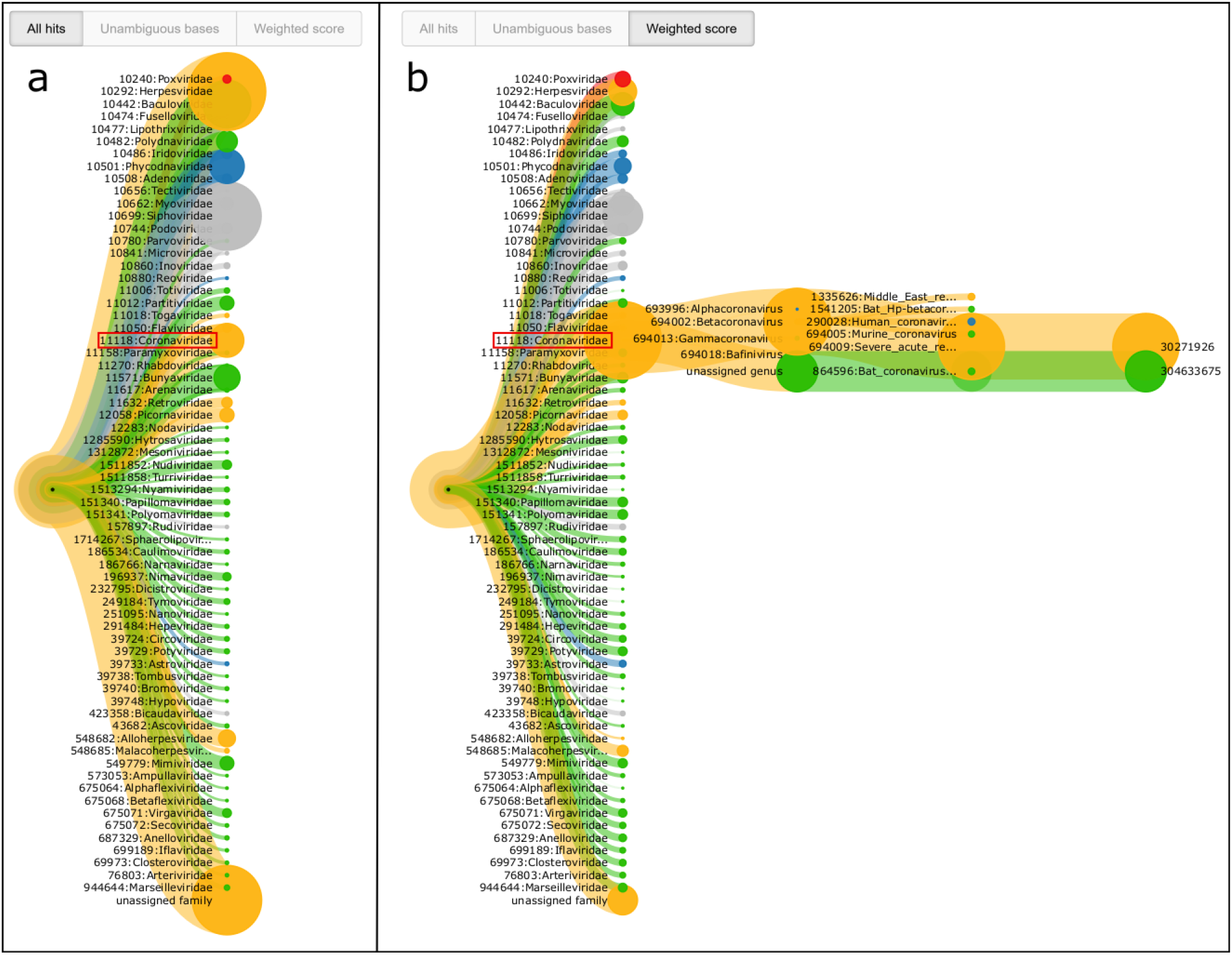
Visualization of the PathoLive results on a real-world dataset from the Wuhan 2019 SARS-CoV-2 outbreak after 30 sequencing cycles. **(a)** Branch thickness determined by the absolute number of aligned reads *Tn.* **(b)** By selecting the weighted score, the results correctly highlight the presence of a Coronavirus by assigning the clearly highest score. When opening the details of the Coronavirus branch, highest similarity can be determined with SARS-related Coronavirus and a Bat Coronavirus. In both subfigures, the family of Coronaviridae is marked by a red box.

When using the pure quantity of alignments for the visualization of results, the Coronaviridae family branch is not among the most prominently visualized hits. In contrast, when activating the weighted score, a clear indication was already available after only 30 sequencing cycles, corresponding to 10% of the complete sequencing run (Fig. 7). The ranks and underlying scores of the visualization shown in Table 1 further support the strength of the weighted score approach. A more detailed analysis of the underlying tree of the Coronaviridae family shows that there are different Coronavirus species with high scores, mainly dominated by a SARS-related Coronavirus and a Bat Coronavirus. This further indicates that a clear assignment to one of the underlying species was not possible. These results are as expected, since the correct species, SARS-CoV-2, was not present in the reference database.

**Table 1.** Comparison of the ranking of the Coronaviridae family for Total Hits and Weighted score. While the Coronaviridae family is not among the most abundant hits, it clearly shows the highest Weighted Score. The total ranking contains 67 families of which the six highest scores are shown.

| Total hits |  | Weighted Score |  |
| --- | --- | --- | --- |
| Family | Hits | Family | Score |
| 1 Herpesviridae | 21,811 | 1 Coronaviridae | 46,274 |
| 2 unassigned family | 17,345 | 2 Siphoviridae | 11,559 |
| 3 Siphoviridae | 16,838 | 3 unassigned family | 6,148 |
| 4 Baculoviridae | 7,996 | 4 Herpesviridae | 5,342 |
| 5 Phycodnaviridae | 4,018 | 5 Myoviridae | 3,775 |
| 6 Coronaviridae | 3,827 | 6 Baculoviridae | 3,423 |
| ... |  | ... |  |

Another branch that is clearly highlighted by its red color belongs to the Poxviridae family. However, a more detailed look into the results shows that the BSL-4 classification originates from a single sub-branch where all mapping positions cluster to only two single peaks (not shown). This is a similar pattern to what we already showed for the occurrence of HIV1 in the previous section (cf. Fig. 6d) and is therefore most probably not of biological interest.

## 4 Discussion

NGS has been shown to be the current state of the art DNA sequencing technology for pathogen detection and makes an increasing impact on the diagnosis of infectious diseases. Although Third Generation Sequencing approaches are also becoming more and more influential, the discovery of lowly abundant pathogens is still problematic due to the relatively low number of reads. Additionally, the comparably low coverage and high error rates still hamper certain types of complex follow-up analyses such as the detection of antimicrobial resistances or the geographical origin of a pathogen. On the other hand, long-read sequencing technology show an immense potential for real-time diagnostics in the future, especially when considering the continuously decreasing error rates, shorter sample preparation times, arising higher throughput devices such as the PromethION, as well as valuable technology-specific features such as the read until functionality for that first attempts have been made to separate microbial reads from host DNA during the sequencing procedure (Edwards, et al., 2019; Loose, et al., 2016). All these aspects considered we assume long-read sequencing technology a valuable complement to NGS-based diagnostics in future with distinct properties and therefore potentially different application areas.

The high turnaround time of NGS-based diagnostics is a major drawback compared to targeted molecular methods. Past efforts to speed up NGS-based diagnostics have been made but often come with significant disadvantages: Quick, et al. (2015) introduced a fast sequencing protocol for Illumina sequencers that allows obtaining results after as little as 6 hours. This speedup is accompanied by lower throughput and lower data quality, making it less suitable for whole genome shotgun sequencing approaches without a priori knowledge. Other approaches aiming at performing analyses of intermediate sequencing data require either a massive reduction of the amount of analyzed reads and/or targets (Stranneheim, et al., 2014) or the application of specialized hardware such as field-programmable gate array technology (FPGA) which is, for example, used for the DRAGEN system (Miller, et al., 2015). Such specialized hardware approaches come with additional costs, either for purchase and infrastructure of local solutions or for the use of a cloud system. At the same time, such approaches provide a low level of flexibility in the analysis and are not algorithmically optimized for working with incomplete data. PathoLive does not require the use of specialized hardware and provides accurate diagnostics results in real-time, illustrated with an easily understandable and interactive visualization. This strongly facilitates to get insights into a clinical sample before the sequencer has finished. Real-time output before the sequencing process of the first read has finished lacks information about multiplex indices, though. Therefore, early results of multiplexed sequencing runs can only be assigned to a specific sample after sequencing of the multiplex-indices. For paired-end sequencing runs, this still means analyses are still possible far before the sequencer ends, and single-end sequencing runs can produce results at the very moment the indices have been sequenced. A possible solution for this problem is to sequence the indices before the first read, which can pose addressable challenges for cluster identification. As a working solution, many sequencing devices allow paired-end sequencing with different lengths for the first and second reads. It is thereby possible to sequence only a short fragment of the first read to get early access to the multiplex indices. Thus, this approach can be used to obtain de facto single-end reads (i.e., the full second read) while having the multiplex information available from the beginning of the read.

For pathogen identification, we changed the basis for the selection of clinically relevant hits from pure abundance or coverage-based measures towards a metric that takes information on the singularity of a detected pathogen into account. Still, we decided not to completely trust the algorithmic evaluation alone, but provide all available information to the user in an intuitive interactive taxonomic tree. While we assume that this form of presentation allows users to come to the right conclusions very quickly, more sophisticated methods for the abundance estimation especially on strain level exist. Implementing an additional abundance estimation approach comparable to the read reassignment of Clinical Pathoscope (Byrd, et al., 2014) or the abundance estimation of Bracken (Lu, et al., 2017) could enable more accurate results, albeit this would not be applicable trivially to the overall conception of PathoLive.

The sensitivity and specificity of PathoLive varies with the time of a sequencing run. In the beginning, when only little sequence information is available, only a small number of nucleotides specify a candidate hit, leading to comparably high false positive rates. At the end of a sequencing run, the number of sequence mismatches in the longer alignments may lead to the erroneous exclusion of hits, especially when sequencing quality decreases. However, this behavior is implicitly considered by the HiLive2 algorithm which allows for an increasing number of mismatching nucleotides with increasing length of the reads. Still, the results can vary over runtime with the optimal outcome being measured at intermediate cycles if the selected parameters are not well-suited for the specific sample or if the sequencing quality decreases stronger than usual.

Besides these challenges which are unique to PathoLive, similar problems as conventional approaches occur. First, the definition of meaningful reference databases is difficult. No reference database can ever be exhaustive since not all existing organisms have been sequenced yet. Besides that, there may be erroneous information in the reference databases due to sequencing artefacts, contaminations or false taxonomic assignment. The definition of the hazardousness was especially complicated, as to our knowledge no well-established solution for the automated assignment of this information exists. Therefore, the basis for our BSL-levelling approach might not be exhaustive, leading to underestimated danger levels of pathogens that are missing in the underlying BSL list. Furthermore, in-house contaminations, some of which are known to be carried over from run to run on the sequencer while others may come from the lab, could interfere with the result interpretation of a sequencing run. Especially since no indices are sequenced for the first results of PathoLive, comparably large numbers of carry-over contaminations might lead to false conclusions. Candidate contaminations should therefore be kept in mind when interpreting results.

Using in-house generated spiked human plasma samples, we were able to show the advantages of PathoLive not only concerning its unprecedented runtime but also the selection of relevant pathogens. We further show the high sensitivity of our approach by identifying CCHFV in a real sample from Sudan based on a few dozens of reads. While being very fast and accurate, a limitation of PathoLive lies in the discovery of yet unknown pathogens. This is due to the limited sensitivity of alignment-based methods in general, which hampers the correct assignment of highly deviant sequences. However, the analysis of a dataset from the SARS-CoV-2 outbreak in 2019 clearly shows that the detection of novel species that are related to known pathogens is still possible.

Concluding, PathoLive is a helpful tool for accurate and yet rapid detection of pathogens in clinical NGS datasets. The key advantages are the real-time availability of analysis results as well as the intuitive and interactive visualization with down-prioritization of likely irrelevant candidates.

## Acknowledgements

We gratefully acknowledge the support of Claudia Kohl concerning the selection of appropriate datasets. We thank Andrea Thürmer and Aleksandar Radonić for sharing their expertise in Illumina sequencing. We further thank all HiLive contributors for their work on the real-time read mapping approach.

## References

Andrusch, A., et al. PAIPline: pathogen identification in metagenomic and clinical next generation sequencing samples. Bioinformatics 2018;34(17):i715–i721.

Biosafety and Biotechnology Unit. Belgian classifications for micro-organisms based on their biological risks - Definitions. In. https://my.absa.org/Riskgroups; 2008.

Bolger, A.M., Lohse, M. and Usadel, B. Trimmomatic: a flexible trimmer for Illumina sequence data. Bioinformatics 2014;30(15):2114–2120.

Bostock, M., Ogievetsky, V. and Heer, J. D(3): Data-Driven Documents. IEEE Trans Vis Comput Graph 2011;17(12):2301–2309.

Bray, N.L., et al. Near-optimal probabilistic RNA-seq quantification. Nat Biotechnol 2016;34(5):525–527.

Breitwieser, F.P., Lu, J. and Salzberg, S.L. A review of methods and databases for metagenomic classification and assembly. Brief Bioinform 2017.

Breitwieser, F.P., Pardo, C.A. and Salzberg, S.L. Re-analysis of metagenomic sequences from acute flaccid myelitis patients reveals alternatives to enterovirus D68 infection. F1000Res 2015;4:180.

Breitwieser, F.P., et al. Human contamination in bacterial genomes has created thousands of spurious proteins. Genome Res 2019;29(6):954–960.

Brister, J.R., et al. NCBI viral genomes resource. Nucleic Acids Res 2015;43(Database issue):D571–577.

Byrd, A.L., et al. Clinical PathoScope: rapid alignment and filtration for accurate pathogen identification in clinical samples using unassembled sequencing data. BMC Bioinformatics 2014;15:262.

Bzhalava, D., et al. Unbiased approach for virus detection in skin lesions. PLoS One 2013;8(6):e65953.

Cao, M.D., et al. Streaming algorithms for identification of pathogens and antibiotic resistance potential from real-time MinION(TM) sequencing. Gigascience 2016;5(1):32.

Dadi, T.H., et al. SLIMM: species level identification of microorganisms from metagenomes. PeerJ 2017;5:e3138.

Dutilh, B.E., et al. Editorial: Virus Discovery by Metagenomics: The (Im)possibilities. Front Microbiol 2017;8:1710.

Dutilh, B.E., et al. Reference-independent comparative metagenomics using cross-assembly: crAss. Bioinformatics 2012;28(24):3225–3231.

Edwards, H.S., et al. Real-Time Selective Sequencing with RUBRIC: Read Until with Basecall and Reference-Informed Criteria. Scientific Reports 2019;9(1):11475.

Flygare, S., et al. Taxonomer: an interactive metagenomics analysis portal for universal pathogen detection and host mRNA expression profiling. Genome Biol 2016;17(1):111.

Francis, O.E., et al. Pathoscope: species identification and strain attribution with unassembled sequencing data. Genome Res 2013;23(10):1721–1729.

Frey, K.G., et al. Comparison of three next-generation sequencing platforms for metagenomic sequencing and identification of pathogens in blood. BMC Genomics 2014;15:96.

Greninger, A.L., et al. Rapid metagenomic identification of viral pathogens in clinical samples by real-time nanopore sequencing analysis. Genome Med 2015;7:99.

Greninger, A.L., et al. Rapid Metagenomic Next-Generation Sequencing during an Investigation of Hospital-Acquired Human Parainfluenza Virus 3 Infections. J Clin Microbiol 2017;55(1):177–182.

Hong, C., et al. PathoScope 2.0: a complete computational framework for strain identification in environmental or clinical sequencing samples. Microbiome 2014;2:33.

Huson, D.H., et al. MEGAN Community Edition - Interactive Exploration and Analysis of Large-Scale Microbiome Sequencing Data. PLoS Comput Biol 2016;12(6):e1004957.

Kohl, C., et al. Crimean congo hemorrhagic fever, 2013 and 2014 Sudan. International Journal of Infectious Diseases 2016;53:9.

Kostic, A.D., et al. PathSeq: software to identify or discover microbes by deep sequencing of human tissue. Nat Biotechnol 2011;29(5):393–396.

Langmead, B. and Salzberg, S.L. Fast gapped-read alignment with Bowtie 2. Nat Methods 2012;9(4):357–359.

Lecuit, M. and Eloit, M. The diagnosis of infectious diseases by whole genome next generation sequencing: a new era is opening. Front Cell Infect Microbiol 2014;4:25.

Lecuit, M. and Eloit, M. The potential of whole genome NGS for infectious disease diagnosis. Expert Rev Mol Diagn 2015;15(12):1517–1519.

Lee, A.Y., Lee, C.S. and Van Gelder, R.N. Scalable metagenomics alignment research tool (SMART): a scalable, rapid, and complete search heuristic for the classification of metagenomic sequences from complex sequence populations. BMC Bioinformatics 2016;17:292.

Li, Y., et al. VIP: an integrated pipeline for metagenomics of virus identification and discovery. Sci Rep 2016;6:23774.

Lindner, M.S. and Renard, B.Y. Metagenomic abundance estimation and diagnostic testing on species level. Nucleic Acids Res 2013;41(1):e10.

Lindner, M.S. and Renard, B.Y. Metagenomic profiling of known and unknown microbes with microbeGPS. PLoS One 2015;10(2):e0117711.

Loka, T.P., Tausch, S.H. and Renard, B.Y. Reliable variant calling during runtime of Illumina sequencing. Scientific Reports 2019;9(1):16502.

Loose, M., Malla, S. and Stout, M. Real-time selective sequencing using nanopore technology. Nat Methods 2016;13(9):751–754.

Lu, J., et al. Bracken: estimating species abundance in metagenomics data. PeerJ Computer Science 2017;3:e104.

Menzel, P., Ng, K.L. and Krogh, A. Fast and sensitive taxonomic classification for metagenomics with Kaiju. Nat Commun 2016;7:11257.

Miller, N.A., et al. A 26-hour system of highly sensitive whole genome sequencing for emergency management of genetic diseases. Genome Medicine 2015;7(1):100.

Mokili, J.L., Rohwer, F. and Dutilh, B.E. Metagenomics and future perspectives in virus discovery. Curr Opin Virol 2012;2(1):63–77.

Naccache, S.N., et al. A cloud-compatible bioinformatics pipeline for ultrarapid pathogen identification from next-generation sequencing of clinical samples. Genome Res 2014;24(7):1180–1192.

Norling, M., et al. MetLab: An In Silico Experimental Design, Simulation and Analysis Tool for Viral Metagenomics Studies. PLoS One 2016;11(8):e0160334.

O’Leary, N.A., et al. Reference sequence (RefSeq) database at NCBI: current status, taxonomic expansion, and functional annotation. Nucleic Acids Res 2016;44(D1):D733–745.

Piro, V.C., et al. ganon: precise metagenomics classification against large and up-to-date sets of reference sequences. bioRxiv 2019:406017.

Piro, V.C., Lindner, M.S. and Renard, B.Y. DUDes: a top-down taxonomic profiler for metagenomics. Bioinformatics 2016;32(15):2272–2280.

Piro, V.C., Matschkowski, M. and Renard, B.Y. MetaMeta: integrating metagenome analysis tools to improve taxonomic profiling. Microbiome 2017;5(1):101.

Quick, J., et al. Rapid draft sequencing and real-time nanopore sequencing in a hospital outbreak of Salmonella. Genome Biol 2015;16:114.

Roux, S., et al. Benchmarking viromics: an in silico evaluation of metagenome-enabled estimates of viral community composition and diversity. PeerJ 2017;5:e3817.

Roux, S., et al. Metavir 2: new tools for viral metagenome comparison and assembled virome analysis. BMC Bioinformatics 2014;15:76.

Salzberg, S.L., et al. Next-generation sequencing in neuropathologic diagnosis of infections of the nervous system. Neurol Neuroimmunol Neuroinflamm 2016;3(4):e251.

Scheuch, M., Hoper, D. and Beer, M. RIEMS: a software pipeline for sensitive and comprehensive taxonomic classification of reads from metagenomics datasets. BMC Bioinformatics 2015;16:69.

Skewes-Cox, P., et al. Profile hidden Markov models for the detection of viruses within metagenomic sequence data. PLoS One 2014;9(8):e105067.

Snyder, L.A., et al. Next-generation sequencing--the promise and perils of charting the great microbial unknown. Microb Ecol 2009;57(1):1–3.

Stewart, R.D. and Watson, M. poRe GUIs for parallel and real-time processing of MinION sequence data. Bioinformatics 2017;33(14):2207–2208.

Stranneheim, H., et al. Rapid pulsed whole genome sequencing for comprehensive acute diagnostics of inborn errors of metabolism. BMC Genomics 2014;15(1):1090.

Tausch, S.H., et al. RAMBO-K: Rapid and Sensitive Removal of Background Sequences from Next Generation Sequencing Data. PLoS One 2015;10(9):e0137896.

Tausch, S.H., et al. LiveKraken—real-time metagenomic classification of illumina data. Bioinformatics 2018;34(21):3750–3752.

The 1000 Genomes Project Consortium. A global reference for human genetic variation. Nature 2015;526(7571):68–74.

Truong, D.T., et al. MetaPhlAn2 for enhanced metagenomic taxonomic profiling. Nat Methods 2015;12(10):902–903.

Wommack, K.E., et al. VIROME: a standard operating procedure for analysis of viral metagenome sequences. Stand Genomic Sci 2012;6(3):427–439.

Wood, D.E., Lu, J. and Langmead, B. Improved metagenomic analysis with Kraken 2. Genome Biology 2019;20(1):257.

Wood, D.E. and Salzberg, S.L. Kraken: ultrafast metagenomic sequence classification using exact alignments. Genome Biol 2014;15(3):R46.

Wu, F., et al. A new coronavirus associated with human respiratory disease in China. Nature 2020;579(7798):265–269.

Zhao, G., et al. VirusSeeker, a computational pipeline for virus discovery and virome composition analysis. Virology 2017;503:21–30.

Zheng, Y., et al. VirusDetect: An automated pipeline for efficient virus discovery using deep sequencing of small RNAs. Virology 2017;500:130–138.

