## Supplemental Material for "PathoLive – Real-time pathogen identification from metagenomic Illumina datasets"

#### --- Supplementary Material ---

### 1 Supplementary Fig. 1

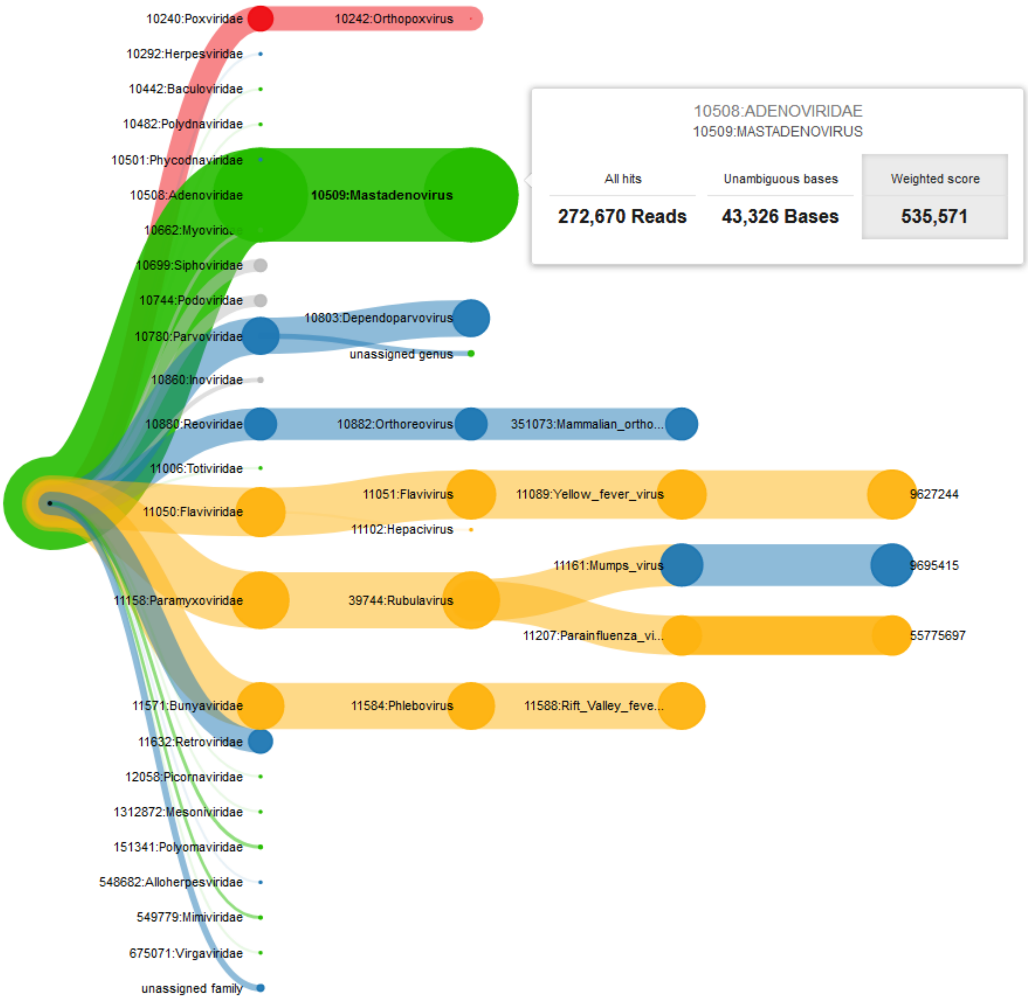

**Supplementary Fig. 1** Example of the interactive taxonomic tree of results of PathoLive for a spiked dataset when using HiLive2 for read alignment.  It shows the visualized results of the described plasma sample at cycle 80 based on the weighted score.

### 2 Supplementary Table 1

**Supplementary Table 1** Comparison of AUC values of PathoLive for a spiked dataset when using HiLive and HiLive2 for read alignment.

| Cycle | AUC value with HiLive | AUC value with HiLive2 |
|---|---|---|
| **40** | 0.94 | 0.94 |
| **60** | 0.92 | 0.92 |
| **80** | 0.92 | 0.92 |
| **100** | 0.90 | 0.88 |

### 3 Supplementary Methods

#### 3.1 Databases used for benchmarking

The reference database for PathoLive was built from the viral part of the NCBI RefSeq downloaded on 2016-07-06. For Clinical Pathoscope we downloaded the associated database from http://www.bu.edu/jlab/wp-assets/databases.tar.gz on 2017-12-09 and used the provided viral database as foreground and the human database as background. The results of Bracken were generated based on the viral part of the NCBI RefSeq downloaded on 2017-12-18. The Bracken database was generated with default parameters and an expected read length of 100 bp.

#### 3.2 Preprocessing of the samples from Sudan and Wuhan

The sequencing data of of the samples from Sudan and Wuhan was only available in preprocessed FASTQ format. To convert the available FASTQ files back to Illumina bcl format, trimmed reads were extended to the required length with calls of the ambiguous nucleotide N. In general, this procedure could influence the results by introducing random hits. However, for the dataset from Sudan we observed that most reads still contained the adapter sequences. As all sequence information after a detected sequencing adapter is ignored from analysis when using the adapter trimming functionality of HiLive2, the applied procedure during format conversion should not have any significant effect on the results in this special case when compared to data directly coming from the sequencing machine. Still, even for reads missing an adapter sequence, as it was the case for the dataset from Wuhan, this would lead to a decrease of aligned reads which further hampers the identification of pathogens and therefore does not limit the validity of the final results. The conversion step from FASTQ to bcl format itself was done by concatenating each read pair in a single FASTQ file and execution of the fastq2bcl script which is delivered with HiLive2. The total length of all reads was 2 x 301bp, corresponding to a total of 602 sequencing cycles, for the dataset from Sudan and 2 x 151bp, corresponding to a total of 302 sequencing cycles, for the dataset from Wuhan.

#### 3.3 Accession numbers of the PathoLive background database

For the creation of the background database we used the datasets from the 1000 Genomes Project Phase 3 with the following accession numbers:

SRR190845, SRR068180, ERR251013, ERR251014, SRR099960, ERR229780, SRR189815, ERR015529, SRR099967, SRR099969, ERR251012, ERR251011, SRR099961, ERR013139, SRR099959, ERR013142, SRR701450, SRR098436, ERR018404, ERR015530, ERR251010, ERR251009, ERR015533, SRR098442, ERR015517, ERR013112, SRR701451, ERR015880, ERR019906, ERR015763, ERR013144, SRR707169, ERR015762, SRR099955, ERR018557, ERR015532, ERR013156, ERR015515, ERR013145, ERR013161, ERR013152, ERR016162, ERR013158, ERR018405, SRR098439, SRR043393, ERR018402, ERR018547, SRR707168, SRR741387, ERR018420, ERR016155, SRR062639, SRR062636, SRR741386, SRR101476, SRR101463, SRR101475, SRR043351, ERR015879, SRR101469, SRR718071, ERR016351, SRR062637, ERR016161, ERR018418, ERR018419, SRR101474, SRR060290, SRR037754, SRR037755, ERR031937, SRR101473, SRR051599, ERR031965, SRR060294, ERR016168, ERR013101, ERR016167, ERR031933, SRR101466, SRR101470, SRR764703, SRR037756, SRR101472, SRR035595, SRR038565, ERR016158, SRR060289, ERR016345, SRR037753, SRR764730, ERR016157, SRR035596, SRR101471, SRR101478, ERR016350, SRR701480, SRR044231, SRR765995, SRR101464, SRR044232, ERR031964, SRR101465, SRR035677, ERR034564, SRR060292, SRR060291, SRR044233, SRR766045, ERR031932, SRR707198, SRR060293, SRR101467, SRR711355, ERR031936, ERR031935, SRR044235, SRR060295, SRR060296, ERR016160, SRR711356, SRR035676, SRR707196, SRR038561, SRR038564, ERR031934, SRR038563, SRR043360, SRR035673, SRR043357, SRR043396, SRR035600, SRR101477, SRR043410, SRR035674, SRR038562, SRR035675, SRR043354, SRR043384, SRR043392, SRR101468, SRR035594, SRR035593, SRR035672, SRR043379, SRR043372, SRR035591, SRR043378, SRR043381, SRR043386, SRR035592, SRR043370, SRR768526, SRR043382, ERR016005, SRR043405, SRR035590, SRR035601, SRR037782, SRR035589, ERR013146, SRR037783, ERR018521, ERR013131, SRR718072, SRR764729, SRR701483, SRR764704, SRR037777, ERR019904, SRR070801, ERR018523, SRR070516, ERR015527, SRR233084, SRR316803, SRR233083, SRR233086, SRR233075, SRR233102, SRR233105, SRR233085, SRR233088, SRR233069, SRR233079, SRR233087, SRR233074, SRR233101, SRR233082, ERR016166, ERR016159, ERR016156, ERR016169, ERR018403, ERR016163, ERR016165, ERR016164, SRR098444, SRR098432, SRR098438, SRR233073, SRR316801, SRR098437, SRR098441, SRR098433, SRR233107, SRR233106, SRR098435, SRR233097, SRR233104, SRR233094, SRR233078, SRR233091, SRR233096, SRR233071, SRR233100, SRR233099, SRR233089, SRR107017, SRR101146, SRR101150, SRR101144, SRR101145, SRR101147, SRR101148, SRR101149, SRR043361, SRR043362, SRR035485, SRR043383, SRR043408, SRR043367, SRR035484, SRR043356
